# Temporal regulation of metabolic processes in the marine diazotroph *Crocosphaera watsonii* WH 8501

**DOI:** 10.1101/2025.07.11.664471

**Authors:** Julia M. Gauglitz, Keisuke Inomura, Wout Bittremieux, Dawn M. Moran, Matthew R. McIlvin, Mak A. Saito

**Affiliations:** Marine Chemistry and Geochemistry Department, Woods Hole Oceanographic Institution (WHOI), Woods Hole, MA, USA; Graduate School of Oceanography, University of Rhode Island, Narragansett, Rhode Island, USA; Department of Computer Science, University of Antwerp, Antwerp, Belgium

**Author notes:** Department of Computer Science, University of Antwerp, Antwerp, Belgium (Julia M. Gauglitz).

## Abstract

Marine diazotrophic cyanobacteria play a crucial role in oceanic nitrogen cycling, supporting primary production and ecosystem balance. *Crocosphaera watsonii* WH8501 exemplifies this ability by temporally separating photosynthesis and diazotrophy to sustain metabolism. To investigate the regulatory mechanisms underlying this process, we employed LC/MS-MS proteomics in a diel culturing experiment, revealing tightly coordinated protein abundance patterns.

Our findings showed a sophisticated temporal regulation of metabolic processes categorized within six distinct protein abundance clusters: (1) nitrogen fixation and amino acid biosynthesis proteins peaked during the night, while (2) glycogen metabolism and dark reactions of photosynthesis were most abundant during the night and day-night transition, likely supporting carbon consumption and energy production. Midday (3 and 4) was dominated by proteins related to photosynthesis, cellular division, and lipid synthesis, whereas late-day peaks (5) in peptide biosynthesis may facilitate nitrogenase complex formation. Notably, the day-night transition (6) exhibited fine-tuned coordination of nitrogenase assembly, with FeS cluster proteins preceding peak nitrogenase iron protein abundance, implying a temporally ordered sequence for functional enzyme formation. Within these categories, sharp temporal patterns emerged in iron trafficking to heme and iron cluster biosynthetic systems, consistent with the need to maintain tight control of iron distribution to metalloproteins at each temporal transition. These results highlight the intricate diel regulation that enables *Crocosphaera* to balance nitrogen fixation and photosynthesis within a single cell. The observed coordination supports the existence of a complex regulatory system ensuring optimal metabolic performance, reinforcing the critical role of temporal control in sustaining these globally significant biological processes.

## Introduction

Marine cyanobacteria are oxygenic photoautotrophs that are responsible for up to 50% of primary production in surface waters^1^. In addition to their importance to carbon flux in microbial food webs, some species are diazotrophic, or able to fix atmospheric N_2_ into a bioavailable form^2^. While cyanobacterial diazotrophs, such as the filamentous non-heterocystous *Trichodesmium*, were long thought to be the primary source of nitrogen fixation, recent insights indicate a prominent role for unicellular marine cyanobacteria such as *Crocosphaera watsonii* WH8501, which have been shown to substantially contribute to N_2_ fixation in tropical and subtropical waters^3–6^.

The dual role of a diazotrophic cyanobacterium brings with it specific challenges as photosynthesis and nitrogen fixation have an incompatibility that may complicate occurrence in the same milieu at the same time. The key enzyme in nitrogen fixation, nitrogenase, relies on an anaerobic environment^7,8^ and will be inactivated by oxygen which evolves from photosystem II during light dependent carbon fixation. Therefore, most microorganisms have evolved to employ either spatial or temporal separation of diazotrophy and photosynthesis. *C. watsonii* WH8501 is able to perform both of these processes by employing temporal separation^9^ and the metabolic shift between inorganic C fixation during the day and N_2_ fixation at night has now been well established^10,11^.

The nitrogenase enzyme is encoded in the nif operon, which contains the three structural proteins of the nitrogenase complex (NifH, NifD, and NifK) as well as other proteins involved in cofactor biosynthesis and enzyme assembly. While gene transcript studies in *C. watsonii* WH8501 have demonstrated a diel pattern whereby the nif operon transcript levels cycle together over the day/night cycle^12^, presence does not guarantee active function as the nitrogenase complex components are require several other proteins to achieve catalytic competency and form the active nitrogenase enzyme^13^. Interestingly, not all of the functions of the proteins necessary for optimal nitrogenase function have been determined, despite the importance of nitrogen fixation for global biogeochemical cycles and agriculture.

Oxygen management and protection of nitrogenase include increasing membrane diffusivity, increasing respiration (respiratory protection), and protecting and repairing nitrogenase^15,16^. A modeling study has suggested that *Crocosphaera* may have low membrane diffusivity against oxygen at least during the dark period^17^, and a genomics study shows that a hopanoid lipid may be responsible for such low diffusivity^18^. During respiratory protection, the level of respiration can be so high that it has been predicted to substantially lower growth efficiency^19^ and growth rate^20^, but the average cost of respiratory protection for *Crocosphaera* may be decreased by colony formation and metabolism specialization^21^. Previous work has suggested that multiple protection mechanisms are in operation simultaneously^17,19^, enabling *Crocosphaera’s* unique physiological capabilities.

A number of behaviors discriminate *C. watsonii* WH8501 from any other phototroph, and likely play a role in facilitating *Crocosphaera*’s adaptation and competition in the oligotrophic open ocean environment. *C. watsonii* has been shown to meet its physiological requirement for iron in part via the temporal separation of nitrogen fixation and photosynthesis, enabling it to recycle iron atoms between key components of these two separate metabolic processes^22^. Furthermore, temporal separation is marked by the cessation of photosystem II activity during the dark phase of the diel cycle^23^ and *C. watsonii* can fix nitrogen, even in the presence of nitrate (NO_3_ ^-^)^24^, indicating an expanded role for nitrogen fixation in the oceans^25^. Notably, *nifH* genes from related cyanobacterial lineages have been detected in oxygen minimum zones^26,27^, where low ambient oxygen concentrations may facilitate nitrogen fixation. As current projections indicate expansion of oxygen minimum zones, cyanobacterial unicellular nitrogen fixers such as *Crocosphaera* may also begin to play a role in generating biological nitrogen in additional regions of the world’s oceans^24^.

Cyanobacteria are known for their ability to exhibit circadian rhythms, which are driven by a circadian clock^26^, via mechanisms such as the oscilloid model established in the model organism *Synechococcus elongatus* PCC 7942. The oscilloid model is based on a three-protein oscillator system called kaiABC, where gene expression is regulated by the phosphorylation state of the kaiABC protein complex, which is embedded in a network of input and output factors^27,28^. A robust clock model consists of *kaiA*, *kaiB*, *kaiC*, *sasA*, *cikA*, and *rpaA*^29^. The *cikA* gene codes for a bacteriophytochrome-like histidine kinase and plays a role in the input signaling of the clock by sensing light via the detection of quinones via its C-terminal pseudo-receiver domain^30^, providing a link between cellular metabolism and the clock. The concentration and redox state of quinones, which are light dependent, may act to entrain the rhythmicity of the circadian clock to environmental stimuli. *rpaA* acts as a master regulator of global gene expression controlled by the circadian clock^31^. *rpaA* is regulated by *cikA* and an antagonistic histidine kinase—sasA^32^. The latest investigations into circadian clocks in *S. elongatus* PCC 7942 have established an *in vitro* model^33^ which demonstrates the key role of output factors in the functioning of the circadian clock.

Protein abundances of potential circadian clock input and output factors have not been reported for *C. watsonii* WH8501, although it exhibits diel cycling of transcripts, contains genes for the kaiABC protein complex, and has been shown to exhibit changes in DNA topology^11^ consistent with the oscilloid model established in *S. elongatus* PCC 7942. Observational studies are required to help determine whether the oscilloid model is the key form of temporal regulation. Diel gene expression experiments frequently capture several timepoints in the dark and light phases or make a binary comparison between the light phase and the dark phase (i.e. noon and midnight)^34,35^. In particular, we hypothesize that more frequent sampling, particularly at the transition from light to dark, may yield insights into the metabolic restructuring and respiratory protection mechanisms particular to this diazotrophic cyanobacterium.

Here we present evidence from a high-resolution diel experiment of axenic *C. watsonii* WH8501. We used liquid chromatography tandem mass spectrometry (LC-MS/MS) proteomic techniques to perform a diel experiment with 1 to 2 hour sampling intervals. This experiment was then also compared to a prior lower resolution diel experiment with the same strain grown under the same conditions^22^, confirming many details, while also uncovering new facets with the higher temporal resolution. Together, both datasets support the finding that the proteins involved in nitrogen fixation have a distinct ordering of peak abundance, shed light on the circadian mechanisms, and generally expand our knowledge of temporal proteome changes over the diel cycle in the globally important cyanobacterial diazotroph *C. watsonii* WH8501.

## Materials and methods

### Culture conditions

*Crocosphaera watsonii* WH8501 was originally isolated from the tropical/subtropical South Atlantic Ocean^34^ and grows in waters between 26 and 32 °C^35^. Here *C. watsonii* WH8501 was cultivated axenically in acid-cleaned borosilicate glass culture flasks in SO medium^36^. The culture was verified axenic by growth tests in LB and by epifluorescence microscopy (Carl Zeiss, Thornwood, NY, USA). Batch cultures were maintained at 27 °C with a 14:10 light/dark cycle in a diel incubator (Percival). The lighting system mimics the natural daily light cycle by turning on in the morning at a low light intensity then slowly ramping up to a peak intensity at noon before diminishing during the afternoon before turning off for the night. The light intensity ranged from 0 to 120 lum/ft^2^ units.

*C. watsonii* was cultured in an acid-cleaned, autoclaved 20L glass carboy with cotton plug and spigot by addition of 1L of culture to 16L sterile filtered SO media prepared with oligotrophic Sargasso Sea seawater, with no added nitrogen source. Cells were held in suspension and the culture was aerated by pumping sterile air onto the surface of the culture while the culture was slowly stirred with a stir bar. The culture was monitored daily for growth by flow cytometry on a live sample, and two aliquots were fixed with paraformaldehyde and archived.

We began the diel experiment when the culture reached a log phase of growth and samples were collected every 1-2 hours over a 25 hr period (n = 16). Sampling time began at t=0. Light phase is timepoints 0 (0), 1 (1.5hrs), 2 (3hrs), 3 (4.5hrs), 12 (18.5 hrs), 13 (20.5 hrs), 14 (22.5 hrs), 15 (24 hrs) and dark phase is timepoints 4 (5.5hrs), 5 (6.5hrs), 6 (8.5hrs), 7 (10.5), 8 (12.5), 9 (14.5), 10 (15.5), 11 (16.5hrs). Red lights were used to illuminate the work area to minimize the impact of external cues on cellular metabolism during opening and closing of the incubator. During dawn and dusk the sampling was every hour in order to capture the transitions in protein abundances associated with the onset of nitrogen fixation and photosynthesis. Approximately 50mL culture was drained to flush the spigot, after which samples were collected at each time point. Subsamples were taken for cell counts (fixed with paraformaldehyde and run on a Guava easyCyte HT) and nitrogen fixation rate measurements (acetylene reduction assay, below). Cell counts over the course of the experiment can be found in the Supp. Info. (**SI Fig 1**). Samples were centrifuged at 12,428 rcf for 20 minutes at 4°C and the cell pellet was transferred and microcentrifuged at 6,700 rcf for 8 minutes at 4 °C, further decanted and frozen at -80 °C until further analysis.

### Acetylene reduction assay

As a proxy for nitrogen fixation, the acetylene reduction assay was performed by injecting 2 mL of concentrated acetylene gas into the head-space of a sealed 60 mL Nalgene culture bottle containing 20 mL of the sampled culture located in the incubator. The culture was incubated for one hour after which the headspace was measured for the reduction product (ethylene) by gas chromatography on a Shimadzu GC-8A using hydrogen as carrier gas and calibrated to a 9.1 ppb ethylene standard. One or two samples were collected at timepoints 3–9 to confirm active nitrogen fixation (**SI Fig 2**).

### Protein digestion and global proteomic analysis

Proteins were analyzed by a data-dependent acquisition LC-MS/MS method. Proteins were extracted by a magnetic bead method (Sera-Mag SpeedBeads (GE Healthcare Life Sciences)) and trypsin digested using a modification of previously published methods^22,37^ as described in Gauglitz et al. (2021)^38^. Protein abundance was quantified by a colorimetric BCA protein assay (Thermo Fisher), and peptide extract in 2% DMSO was quantified using a colorimetric assay at 750 nm (DC protein assay; Biorad). Peptides were acidified with 1% formic acid and brought to a final concentration of 0.1 µg/uL with an LC-MS grade buffer containing 98% water, 2% ACN, and 0.1% formic acid.

Acidified protein extracts were analyzed by LC-MS/MS using a Michrom Advance HPLC coupled to a Thermo Scientific Orbitrap Fusion mass spectrometer with a Michrom Advance CaptiveSpray source. Each sample (1µg protein measured before tryptic digestion) was concentrated onto a trap column (0.2 x 10mm, 5µm particle size, 120 Å pore size, ReproSil-Gold, Dr. Maisch GmbH) and rinsed with 100µL 0.1% formic acid, 5% ACN, 95% water before gradient elution through a reverse phase C18 column (0.1 x 250mm, 3µm particle size, 120 Å pore size, ReproSil-Gold, Dr. Maisch GmbH) at a flow rate of 0.25 µL/minute. The chromatography consisted of a nonlinear 210 min gradient from 5% to 95% buffer B, where A was 0.1% aqueous formic acid and B was 0.1% formic acid in ACN (all solvents were Fisher Optima grade). The mass spectrometer was set to perform MS/MS on the top N ions (Thermo Fusion) using data-dependent settings (dynamic exclusion 15s, excluding unassigned and singly charged ions). Ions were monitored over a range of *m/z* 380–1580.

### Data processing

Mass spectral data (16 raw files) were processed in Proteome Discoverer 1.4 using SEQUEST (version 1.4.1.14) and Scaffold 4.4.6 (Proteome Software). Peptide identifications were made implementing experiment-wide grouping with binary peptide-protein weights, a 97.0% peptide threshold and a 99.0% minimum protein threshold with 2 peptides minimum at a 1.6% protein false discovery rate (FDR) and 0.0% peptide FDR. Mass spectra were mapped to the *Crocosphaera watsonii* WH8501 proteome (version November 2008; 5659 entries; **SI File 1**) using the following search parameters: 0.60 Da fragment tolerance; 10.0 ppm parent mass tolerance, fixed modification of +57 on C (carbamidomethyl), variable modification of +16 on M (oxidation) and 2 max missed cleavages. A total of 1148 proteins were identified, which represents a genome coverage of 20%.

All proteomics data are given as relative protein abundance based on normalized spectral counts using the NSAF (normalized spectral count abundance factor) method implemented in Scaffold 4.4.6 (Proteome Software). Fold changes between light and dark phase were calculated as follows: mean values were computed for the light phase and dark phase separately, while imputing zero values by half of the minimum measured abundance across all proteins. P-values were calculated using the T-test based on all abundance values between the light phase and the dark phases, and the Benjamini-Hochberg procedure was used for multiple testing correction.

### Protein clustering

Protein abundances across all timepoints were analyzed using agglomerative clustering. First, proteins for which no molecular weight was assigned were removed. Next, the protein abundances were standardized by removing the mean and scaling to unit variance for each individual protein. Finally, the standardized protein abundances were clustered by agglomerative clustering with Ward linkage and the Euclidean distance. The distance threshold was set to 30 after visual inspection of the dendrogram (**Fig 2A**). Data processing employed Pandas (version 2.0.3)^39^, NumPy (version 1.25.2)^40^, SciPy (version 1.11.1)^41^, and Scikit-Learn (version 1.3.0)^42^ for scientific data processing, and matplotlib (version 3.7.2)^43^ and Seaborn (version 0.12.2)^44^ for data visualization. Jupyter notebooks^45^ were used for exploratory data analysis.

### GO analysis

Gene ontology (GO) enrichment analysis for each cluster detected using agglomerative clustering was performed using the find_enrichment.py script from GOATOOLS^46^. For each cluster, the protein identifiers for the proteins in that cluster were used as the study identifiers, all *C. watsonii* WH8501 protein identifiers read from the FASTA file were used as the population identifiers, and an association file mapping between Uniprot identifiers and GO terms retrieved from QuickGO^47^ was used. Computed p-values were corrected using the Benjamini-Hochberg procedure and thresholded at a p-value of 0.05 after multiple hypothesis testing correction. Other configuration options were left at their default values. GO terms are hierarchical short phrases that describe the known biological function within the groups “biological process”, “cellular compartment”, and “molecular function.” The max depth (MD) of GO terms refers to the depth within the hierarchy, with higher numbers indicating a more specialized role within the groups **(SI File 3)**.

The NIH tool protein BLAST (https://blast.ncbi.nlm.nih.gov/Blast.cgi) was used to search for additional protein identifications, particularly for proteins which were annotated as protein of unknown function.

## Results

We have identified proteins with key roles in oxygen management, circadian rhythm, and nitrogen fixation, many of which display changes in abundance explainable by time of day and the dominant metabolic functions in the cell. Of the 1148 proteins identified by LC-MS/MS analysis, ∼19% have a 2-fold or greater change in abundance across light and dark phases (fold change of < -0.5 or > 2; 218/1148). From these proteins, 78 are statistically different between the light and dark phase and have a 2-fold or greater change in abundance between photo periods (**Fig 1**; p-value <0.05, T-test with Benajmini-Hochberg correction).

This study reveals broad scale clustering of proteins based on temporal signatures, reaffirming the key expression patterns observed in prior *C. watsonii* diel experiments^22,48,49^. Six data-driven protein clusters are identified from the diel proteome by agglomerative clustering (**Fig 2**), with greater similarity between clusters 1 and 2 (night) and the clusters 3, 4 (day) and 5, 6 (dawn/dusk). The significant proteins in **Fig 1** (p-value <0.05) are colored by cluster, as defined in **Fig 2**, to visualize significant signatures from the different clusters within the volcano plot output in **Fig 1**. Furthermore, gene ontology enrichment analysis highlights the biological processes which dominate these protein groupings and their temporal trends, such as photosynthesis enriched in the day clusters (cluster 3, 4) and nitrogen fixation and related GO terms enriched for in the night cluster 1 (**Table 1; SI File 3**).

**Figure 1.**
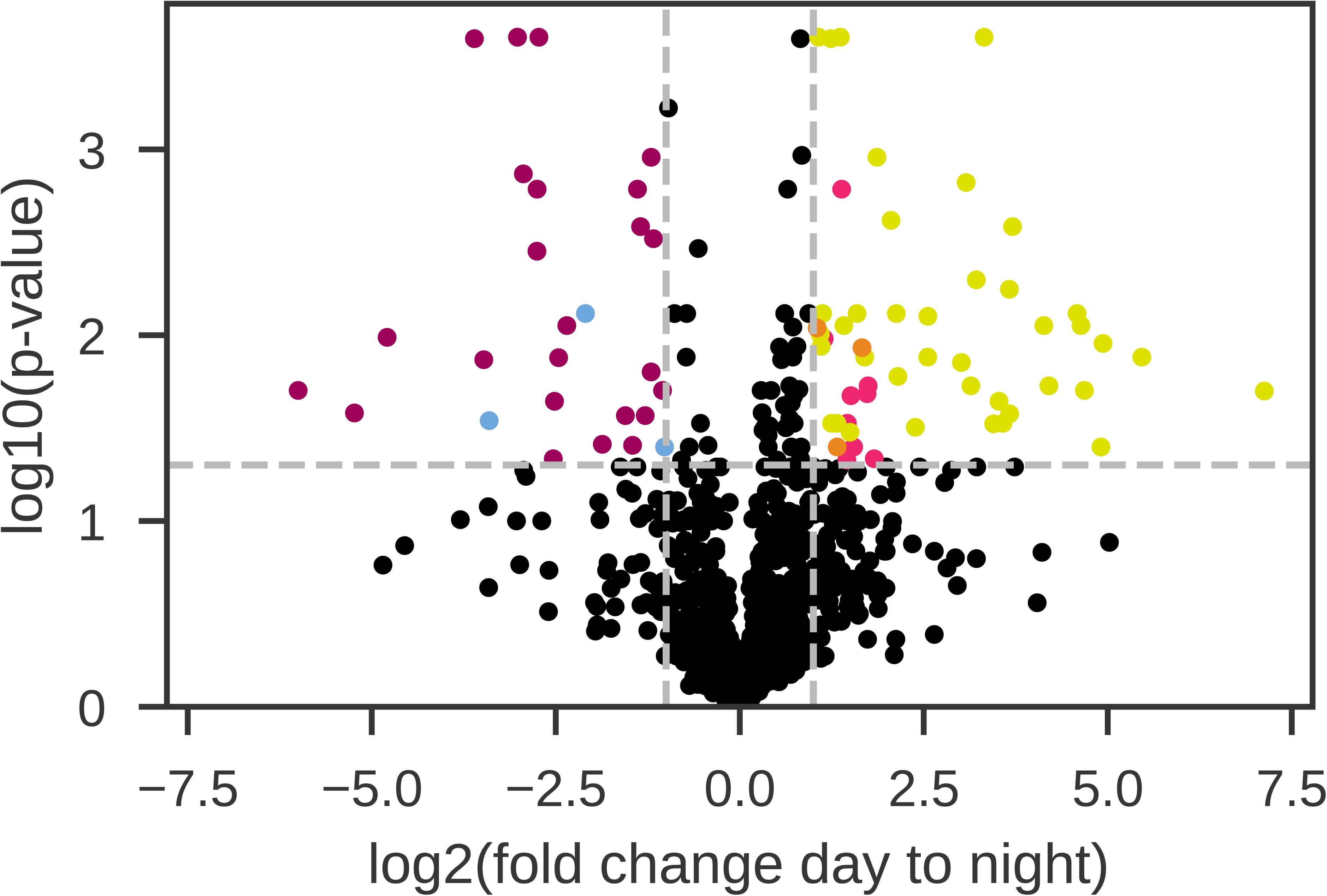
Volcano plot of the global proteome, showing the log2 fold change day to night vs the log10 of the p-value. 27 proteins have statistically higher abundance in the dark period [upper left], while 51 proteins have statistically higher abundance during the light period [upper right]. Proteins that are statistically significant and have a 2-fold or greater change in abundance are colored by cluster based on agglomerative clustering, as seen in Fig 2. [cluster 1: magenta; cluster 2: blue; cluster 3: pink; cluster 4: yellow and cluster 5: orange. No Cluster 6 proteins meet the criteria and thus green is not observed.] Data can be found in **SI Table 1**.

**Figure 2.**
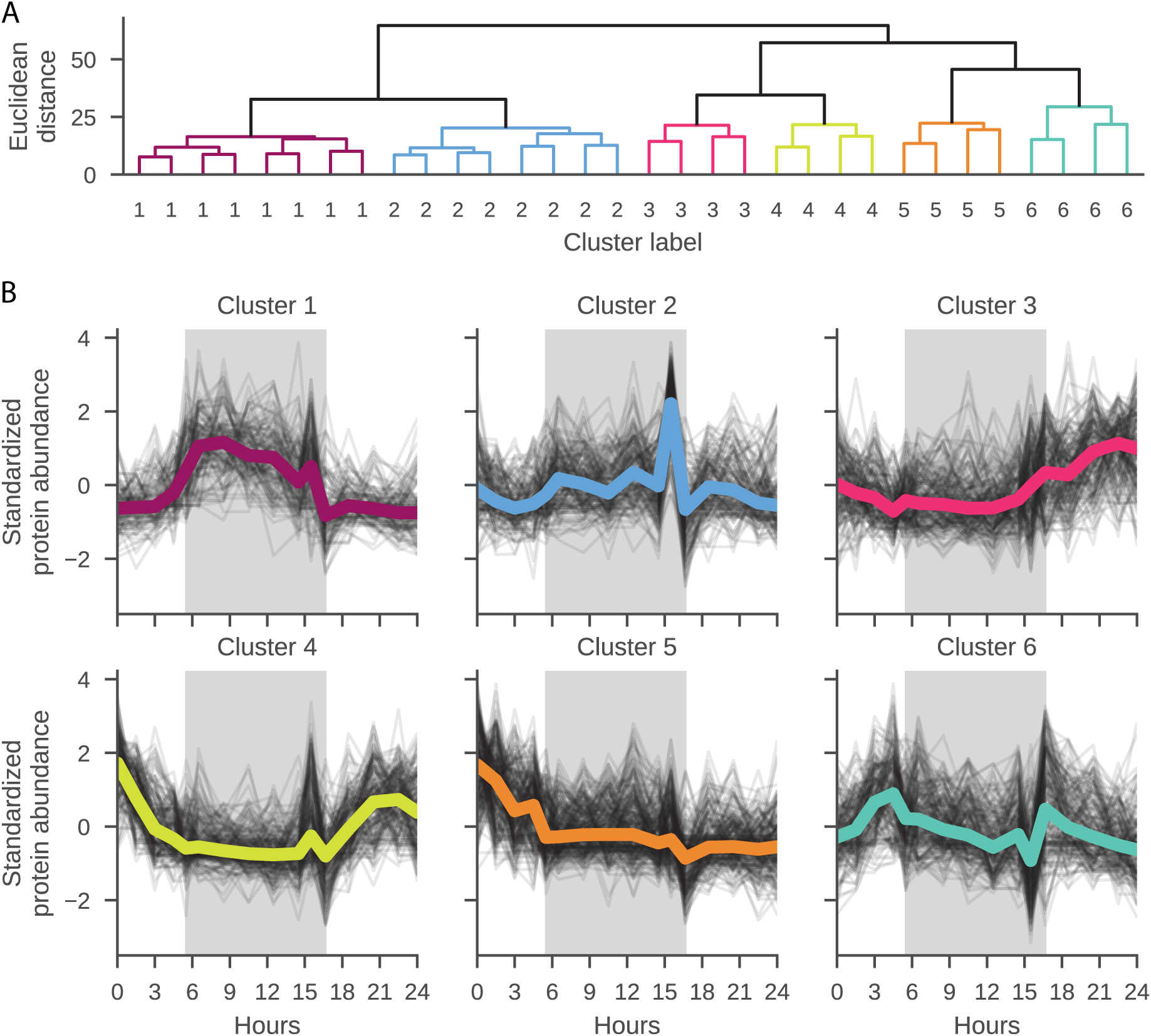
**A.** Proteins clustered by agglomerative clustering with Ward linkage and the Euclidean distance. At a distance threshold of 30 there are six distinct clusters observed. Clusters 1 and 2 have the highest dissimilarity from clusters 3 through 6. **B.** Standardized abundance of proteins grouped with each of the six data-driven clusters are plotted against hours. The dark phase is visualized by the gray background. Average abundance for all proteins in a cluster is colored by cluster as follows: cluster 1: magenta; cluster 2: blue; cluster 3: pink; cluster 4: yellow; cluster 5: orange; cluster 6: green.

**Table 1.** Overview of temporal trends and associated functions by cluster.

| Cluster | Time of max protein abundance | Example systems | Example proteins |
| --- | --- | --- | --- |
| 1 | Night | Nitrogen fixation<br>Amino acid biosynthesis | NifH, NifD, NifK, NifU, NifJ<br>Threonine synthase |
| 2 | Night / Interface | Glycogen metabolic process<br>Photosynthesis, Dark reaction | Glucose-1-phosphate<br>adenylyltransferase,<br>Glycogen synthase<br>Light-independent<br>protochlorophyllide<br>reductase subunit B |
| 3 | Early day | Photosynthesis<br>Carbon fixation | ATP synthase subunits $\alpha$,<br>$\beta$, $\delta$, $\epsilon$<br>Carboxysome shell protein<br>CcmK |
| 4 | Early / Late day | Photosynthesis<br>Cellular division<br>Lipid biosynthesis | Photosystem II<br>MurA, MurE, MurF, Ddl<br>Squalene synthase |
| 5 | Late day | Peptide biosynthesis | Small ribosomal subunit<br>protein uS7<br>Ala/Ser/Val/Gly/Cys/Met/Lys--tRNA ligases |
| 6 | Interface | Oxidoreductase complex<br>Translation | NADH ubiquinone<br>oxidoreductase<br>uS19, S11, L5, S21, L12 |

As expected, the processes enriched in the light phase are predominantly associated with photosynthesis and cellular division, and this concurs with the cluster 4 proteins (yellow) in **Fig 1**, such as Ketose-bisphosphate aldolase (Cwat3318), Photosystem II 12 kDa extrinsic (Cwat0626), and S-layer homology region (Cwat6588), which are also observed in the diel results from Saito et al. (2011)^22^. In addition, an array of biosynthetic proteins are significantly enriched, such as glycosyl transferases (Cwat0194, Cwat1204, Cwat3402, and Cwat3972) and we observe an increased abundance of proteins involved in the Entner-Doudoroff pathway (Cwat3088) and the Calvin Cycle (Cwat3318). The gene ontology term ‘regulation of cell shape’ is enriched with increased abundances of MurA (Cwat0821), MurE (Cwat3769), MurF (Cwat4119), and Ddl (Cwat4357) proteins in the morning, peaking around noon, with minimal protein detected during the night, concurring with cellular division typically occurring in the light period. Of these, UDP-N-acetylglucosamine 1-carboxyvinyltransferase (MurA) and D-alanine-D-alanine ligase (Ddl) protein abundances are also statistically significant with a 2-fold or greater increase (cluster 4).

### Temporally regulated translation of nitrogen fixation components

The nitrogen fixation proteins NifH, NifD, NifK, NifW, NifU, NifJ (Cwat3818, Cwat3819, Cwat3820, Cwat3827, Cwat3817, Cwat1801), Flavodoxin (Cwat4291) (**Table 2**), and Heme oxygenase (Cwat2459), which has been hypothesized to release iron from cytochrome heme groups at night for use in the nitrogenase complex^22^, are among the statistically significant proteins that have a 2-fold or greater increase in abundance during the dark period [magenta-cluster 1, **Fig 1; SI Table 1**]. In addition, we observe NAD(P)(+) transhydrogenase (AB-specific) (Cwat1931) and NAD(P) transhydrogenase, beta subunit (Cwat1933), as well as Glucose–6-phosphate dehydrogenase (Cwat3080), an enzyme in the pentose phosphate pathway, which yields NADPH, the reductant required by nitrogenase. We also detect multiple proteins involved in amino acid biosynthesis and nitrogen metabolism (Threonine synthase (Q4C2V3, Cwat4050), 3-isopropylmalate dehydratase large subunit, (Q4BW83, Cwat1322), Glutamate synthase (ferredoxin) (Q4C8E2, Cwat5417), RNA polymerase sigma factor SigA (Q4C7J7, Cwat5428), Chorismate synthase (Q4BZH4, Cwat2085)), whereby the cell effectively integrates newly fixed nitrogen into metabolic building blocks.

**Table 2.**
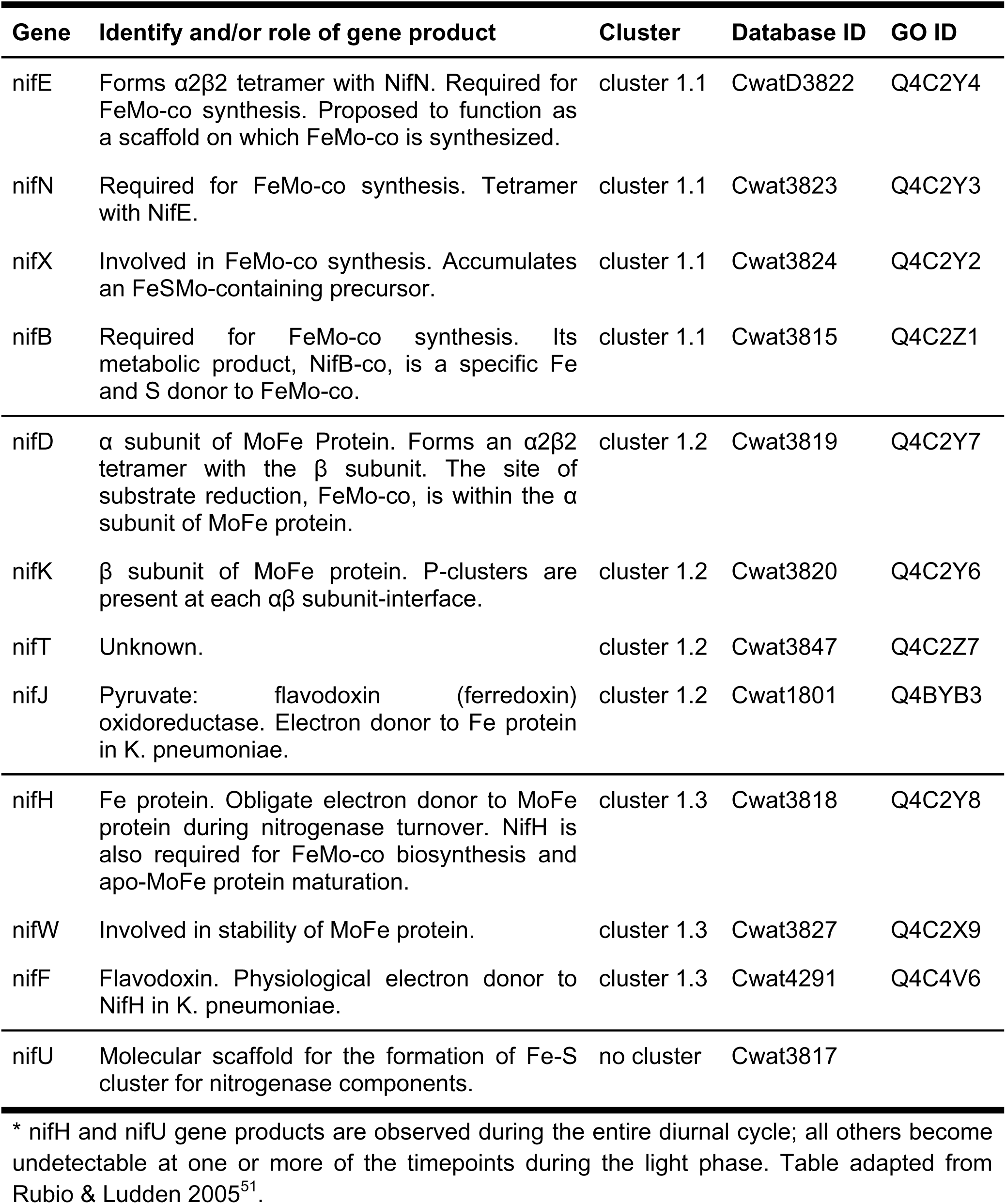
*Nif* gene products, their corresponding protein identity and/or role in nitrogen fixation and clustering pattern. GO ID = UniProt ID.

Nitrogen fixation was the most enriched GO term in cluster 1 with an adjusted p-value of 3.85 e-14, with 11 of 13 proteins enriched for this GO term. We further performed a GO enrichment analysis on only the proteins from cluster 1, employing a lower Euclidean distance cutoff of 11, generating four subclusters, of which cluster 1.4 was not enriched in any terms and is not further discussed (**Figure 3A**). Clusters 1.1, 1.2, and 1.3 contained the enriched term ‘nitrogen fixation’, with a corrected p-value of 1.57 e-4, 1.57 e-4, and 2.28 e-2, respectively. Each of the 11 proteins clustered with one of the subclusters of the dark phase cluster, enabling identification of several nitrogen fixation-related proteins that show maximum relative abundance with different temporal patterns (**Table 2**) in distinct clusters. These results confirm the trends observed in Saito et al.^22^, and due to the higher resolution of this dataset, we are also able to identify further substructure in the abundance patterns of Nif proteins.

**Figure 3.**
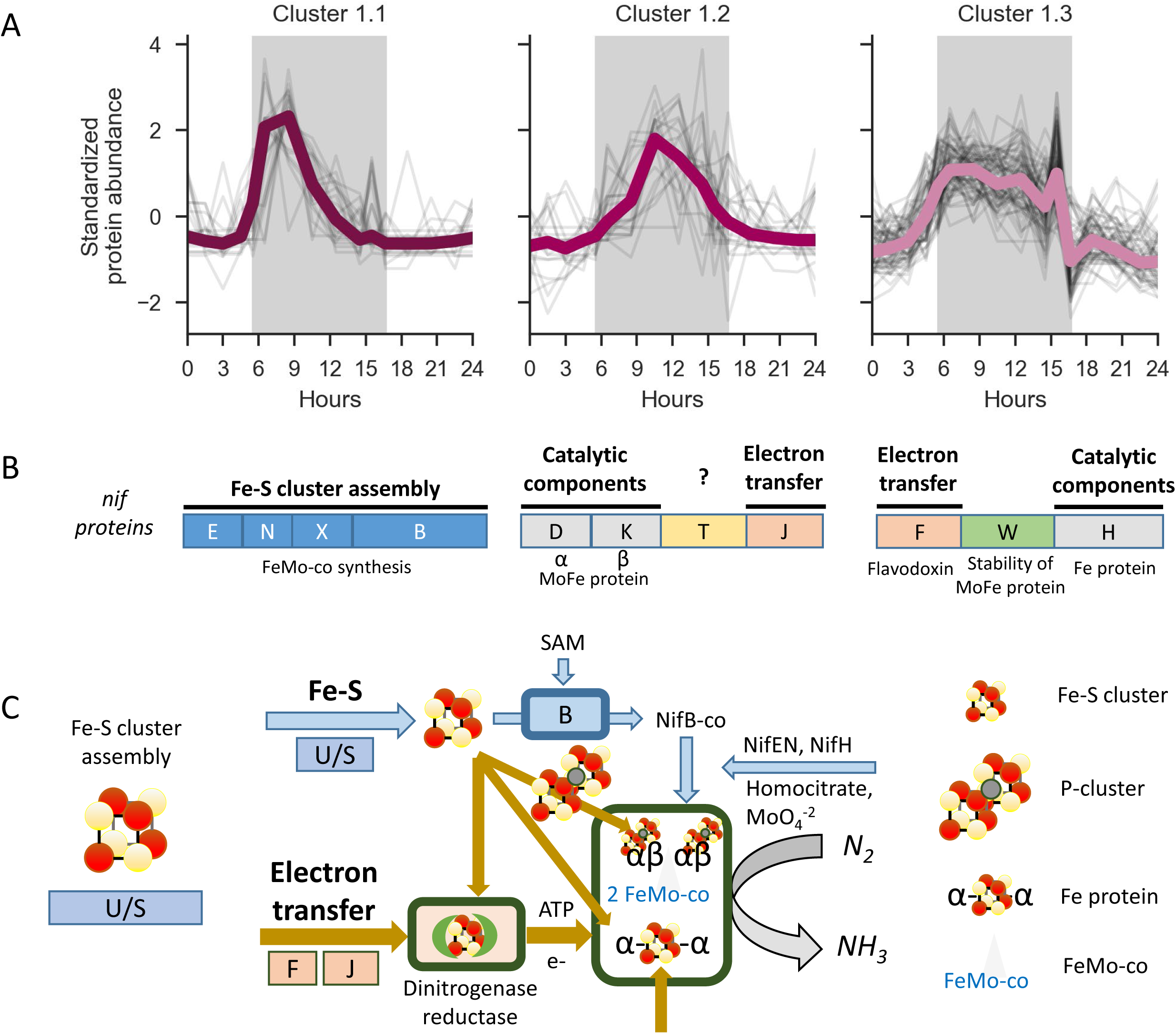
**A.** Temporal progression of Nif biosynthetic groups. Standardized Abundance of proteins grouped with cluster 1 sub-clusters during the diel cycle (cluster 1.4 not shown). The dark phase is visualized by the gray background. Average abundance for all proteins in a cluster is colored by cluster. **B.** nif operon, genes, colors align with schema in part C and are labeled by their role (see also **Table 2**). The genes align with the clusters in A in which they are observed. **C.** Schematic of the role and utilization of the different nitrogenase complex components in dinitrogen fixation. The abbreviations U, S, B, F, J refer to the proteins from the respective nif genes. For example, B is the abbreviation for nifB.

The nitrogen fixation proteins are expressed in a sequential fashion, beginning with a distinct peak in abundance of cofactor biosynthesis proteins (cluster 1.1), followed by a distinct increase in abundance of the MoFe nitrogenase protein subunits (cluster 1.2) (**Fig 3**). In contrast to other gene products necessary for nitrogen fixation, the Nitrogenase iron protein (NifH) is present throughout the dark phase, with maximum Nitrogenase iron protein abundance coinciding with the time of observed maximum nitrogen fixation^12,50^ (**SI Fig 2**), coinciding with the highest abundances of the nitrogenase complex components, such as MoFe nitrogenase protein beta chain (NifK) (**Fig 4**).

**Figure 4.**
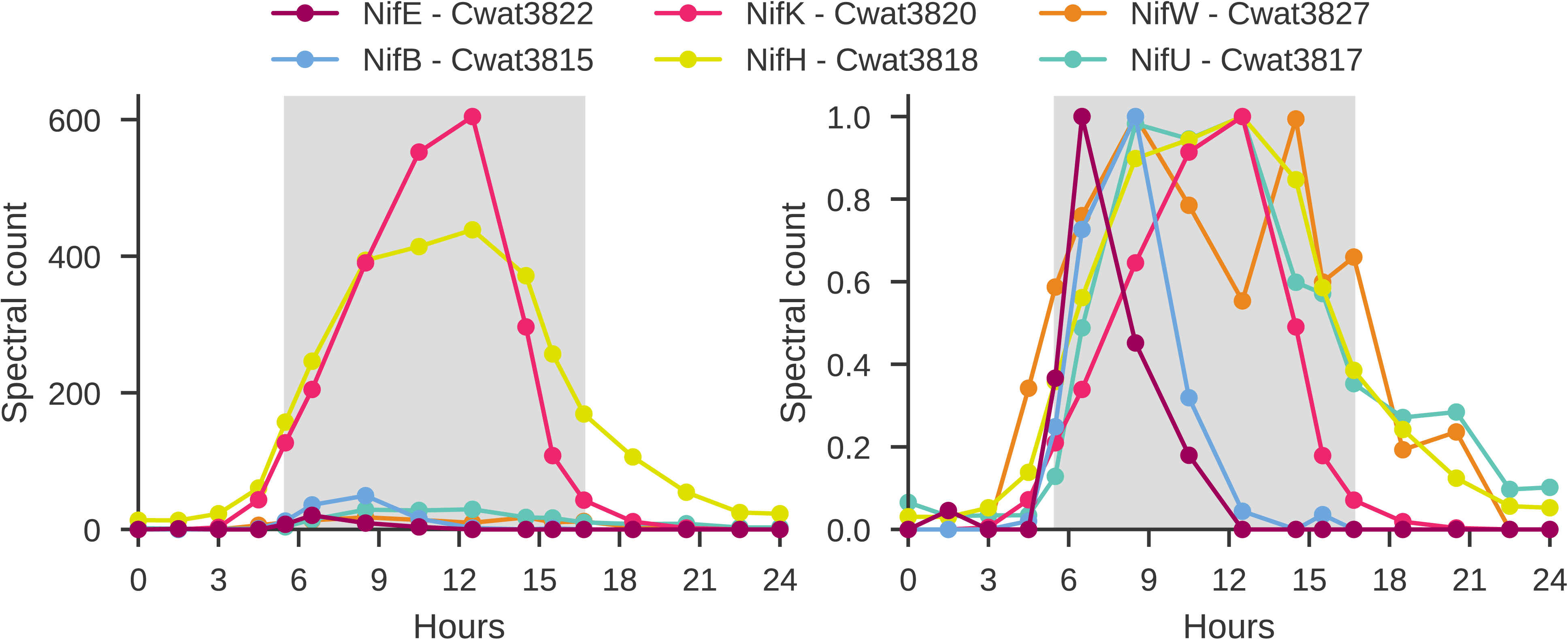
*Crocosphaera* NifB before NifH: Making Fe-S clusters prior to Nitrogenase. MoFe Cofactor biosynthesis protein NifE (Cwat3822 - dark pink); Nitrogen cofactor biosynthesis protein NifB (Cwat3815 - blue); MoFe Nitrogenase protein Beta chain - NifK (Cwat3820 - pink); Iron nitrogenase protein - NifH (Cwat3818 - yellow); Nitrogenase-stabilizing/protective protein NifW (Cwat3827 - orange); Nitrogenase iron-sulfur cluster protein NifU (Cwat3817 - green). Hours are relative to the start of the experiment, with the dark phase visualized by the gray background. **Left.** Spectral counts. **Right.** Spectral counts normalized to peak abundance per protein.

Maturation of the nitrogenase enzyme is a multi-step process and does not occur until the MoFe components are translated. The mature dinitrogenase enzyme is believed to begin with the Fe protein [4Fe−4S] cluster, then the P cluster, and then the FeMo-cofactor, which provides the substrate reduction site (as shown in **Fig 3B, 3C**). NifW binds to apo-MoFe (without FeMo-co), and is thought to stabilize the protein^52^. This order of events is supported by the sequential translation of the proteins from the nif operon (**Fig 4**). We observe that the maximum abundance of NifB (Cwat3815) is four hours prior to that of the Nitrogenase beta chain NifK (Cwat3820). The Fe protein NifH (Cwat3818) has a similar abundance profile with lower levels of Fe protein present earlier in the dark cycle and highest abundance after the halfway mark in the dark cycle. These observations also agree with published reports of peak nitrogen fixation both in culture and in the environment that show the greatest rate of nitrogen fixation co-occurring with peak quantities of the nitrogenase Fe protein^12,50^.

FeS biosynthesis is critical for Nif production. The nitrogenase iron-sulfur cluster protein NifU is involved in assembly and stabilization of the Fe-S clusters in nitrogenase and was observed with higher levels during the night (**Fig 4**). The proteins SufB, SufC, SufD are closely related components of the SUF system, which work together to ensure the proper assembly and maintenance of iron-sulfur clusters in the cell. Their highest abundance is during the first half of the night, although they are present throughout the diel cycle (**SI File 2**)^22^. In addition to biosynthesis, these proteins are also involved in protection of the FeS complexes.

Furthermore, the nitrogenous iron protein (NifH) and NifU are also still detectable at low levels during the light phase. NifU, which acts as a molecular scaffold for the formation of Fe-S clusters for nitrogenase components, may be present at all times to quickly initiate formation of Fe-S clusters, or alternatively, or it may still be observed due to incomplete non-specific protease activity. On the other hand, NifH, the nitrogenous iron protein, is an integral component of the active nitrogenase complex. Although the max abundance of NifH has been shown to correspond with max nitrogen fixation, we note that the presence of NifH does not guarantee active metabolic function. However, it is also required for FeMo cofactor biosynthesis and apo-MoFe protein maturation, which may be the key as to why it is not fully degraded when not active in nitrogen fixation. Furthermore, similar to results from *A. vinelandii*^53^, production of cofactor (nifB) does not require the presence of the nitrogenase protein. We hypothesize that the sequential changes in abundance of these proteins has both to do with metal requirements and metal use as well as maximizing metabolic efficiency, to minimize the need for respiratory protection mechanisms during the early evening while cellular oxygen levels are decreasing.

### Oxygen management and nitrogenase protection

Oxygen management mechanisms may include changes to cell permeability^17,18,54^, controlled redox based conformational change of nitrogenase^55,56^, increased respiration (respiratory protection)^15–17,19^, and physical protection/maintenance of oxygen sensitive proteins^15,56,57^. Prior analyses have postulated that hopanoid lipid synthesis may protect cells from O_2_ by decreasing their permeability^18^. Squalene synthase, a key enzyme in the biosynthesis of hopanoids, displays diel changes in abundance (**Fig 5A**), with a peak during dusk, indicating preparatory synthesis and a key temporal role (cluster 4). In some cyanobacteria, Ni-dependent hydrogenases can help mitigate oxidative stress by consuming excess hydrogen produced during nitrogen fixation, which might otherwise react with oxygen^36^. We observe relatively high abundances (max 52 spectral counts) and entrainment of the Ni-dependent hydrogenase (**Fig 5C**; cluster 1), with abundance patterns aligning with the metabolic demands and environmental conditions encountered at dusk and night. A search in the proteome for other Ni related proteins using protein BLAST against proteins annotated as hypothetical proteins resulted in the identification of a putative Ni transport protein (Cwat4537) expressed only at dusk, prior to the peak of the Ni-dependent hydrogenase (**Fig 5C, blue**). Additional proteins which may participate in broader protective mechanisms that mitigate oxidative stress during nitrogen fixation are radical SAM enzymes, which play an essential role in the biosynthesis and maintenance of the nitrogenase enzyme and its cofactors. Specifically, radical SAM supports the assembly and repair of Fe-S clusters, and its abundance patterns (**Fig 5B**; cluster 4) support an important role in the transition from light to dark.

**Figure 5.**
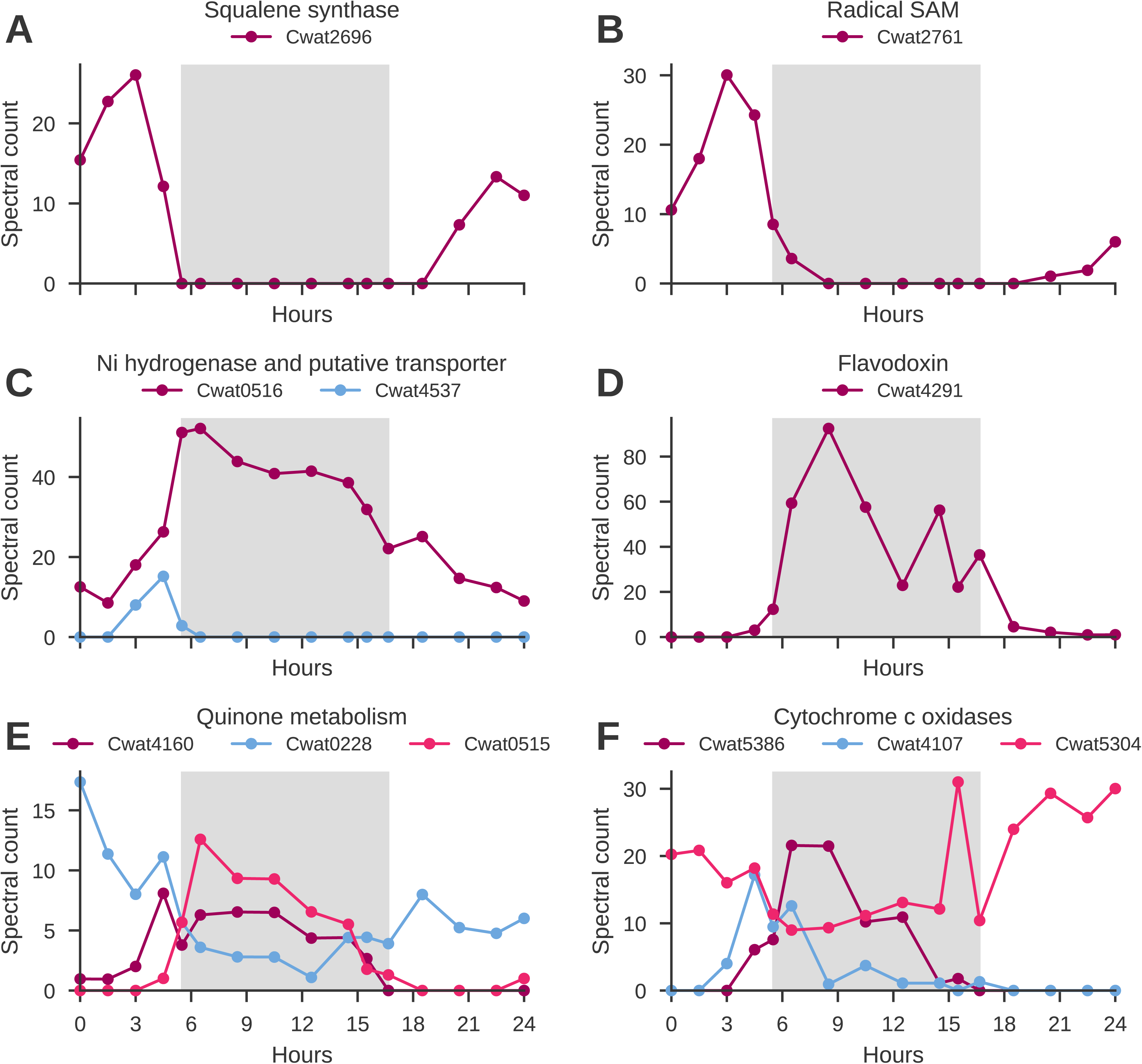
Temporal variability of selected proteins. **A.** Squalene synthase (Cwat2696); **B.** Radical SAM (Cwat2761); **C.** Ni hydrogenase (Q4BUZ7, Cwat0516; purple) and putative Ni transport protein (Cwat4537; blue); **D.** Flavodoxin (Q4C4V6, Cwat4291); **E.** Quinone metabolism (Cwat4160, NADH-plastoquinone oxidoreductase, chain 5, purple); (Cwat0228, similar to Methylase involved in ubiquinone/menaquinone biosynthesis, blue); (Cwat0515, NADH ubiquinone oxidoreductase, 20 kDa subunit, (pink); **F.** Cytochrome c oxidases: (Cwat5386, purple); (Cwat4107, blue); (Cwat5304, pink). The dark phase is visualized by the gray background.

The redox state of the cell may play a role in both entraining circadian rhythms and enabling a favorable environment for nitrogen fixation. We observe temporal patterns for a number of proteins with roles in managing oxidative stress and maintaining the cellular redox balance, including quinones, cytochromes c, and flavodoxin (**Fig 5**). Flavodoxin is more stable under oxidative conditions compared to some other electron carriers, allowing it to effectively participate in redox reactions without being readily inactivated by oxidative damage. It is also used to reduce iron demand in many organisms, but is obligately tied to nitrogen fixation in *Crocosphaera*. Although the primary function of cytochromes is electron transport, we postulate that the increase in cytochromes Cwat4107 and Cwat5386 (**Fig 5F**) may be related to respiratory protection, as increased respiration would result in faster removal of oxygen from the cellular environment.

### Circadian clock in *C. watsonii*

We observe evidence of the oscilloid model of circadian rhythm in the proteome of *C. watsonii* WH8501. Diel cycling of proteins has been reported^22^ and nifH transcript cycling and topological changes seen in DNA staining both support the oscilloid model^11^. We further draw our comparisons of proteins observed to genetic studies in other prokaryotes, primarily from literature on the model cyanobacterium *Synechococcus elongatus* PCC 7942, for which the oscilloid model is described in the introduction above.

The kai genes central to the oscilloid model are present in the *C. watsonii* WH8501 genome, with one copy of kaiA and kaiB and two copies of kaiC, of which one homolog of each gene is together in one gene neighborhood (KaiA - Cwat4942, KaiB - Cwat4943, KaiC - Cwat4944, and KaiC - Cwat5154). These proteins, while detected, do not exhibit a clear temporal trend (**SI Figure 3**). This is, however, not unexpected. As KaiC phosphorylates and dephosphorylates rhythmically during the course of a day, no stark changes in protein abundance are expected, but rather a change in phosphorylation state^26^, which were not measured in the course of this study.

As the core clock proteins are furthermore embedded in a network of input and output factors, we searched for additional signatures in the proteome. While sasA and ldpA homologs are present in the genome based on BLAST alignment, we did not detect the corresponding proteins in the current proteome analysis. In addition, no homologous proteins were identified in the genome used for this analysis for CikA, Pex, or CpmA, however multiple proteins putatively annotated as containing TPR repeats (a structural motif consisting of a degenerate 34 amino acid tandem repeat) are observed with diel patterns and TPR motifs are a key structural component of CikA and are known to play a crucial role in its function (**SI Table 1**, higher abundance during the day; cluster 4&5). These motifs enable CikA to interact with other components of the circadian clock, thereby facilitating the transmission of environmental signals to the clock machinery^58^. Additional potential circadian input/output factors present in the proteome are putative proteins LabA and LalA (based on putative assignment to proteins of unknown function based on similarity searches), and RpaB (Cwat5773; Q4C818), although they do not exhibit oscillating abundance patterns.

The response regulator RpaA (Response regulator receiver:Transcriptional regulatory protein, C-terminal, with 98.4% amino acid sequence identity to RpaA using BLASTp (Cwat6453; Q4C9J3)) contributes to the enrichment in cluster 4 [biological process; molecular function; binding]. This putative RpaA regulatory protein displays a clear diel pattern (**Fig 6A**), which was also observed in the diel data from Saito et al.,^22^. RpaA is a member of a two component regulatory system and activated by phosphorylation.

**Figure 6.**
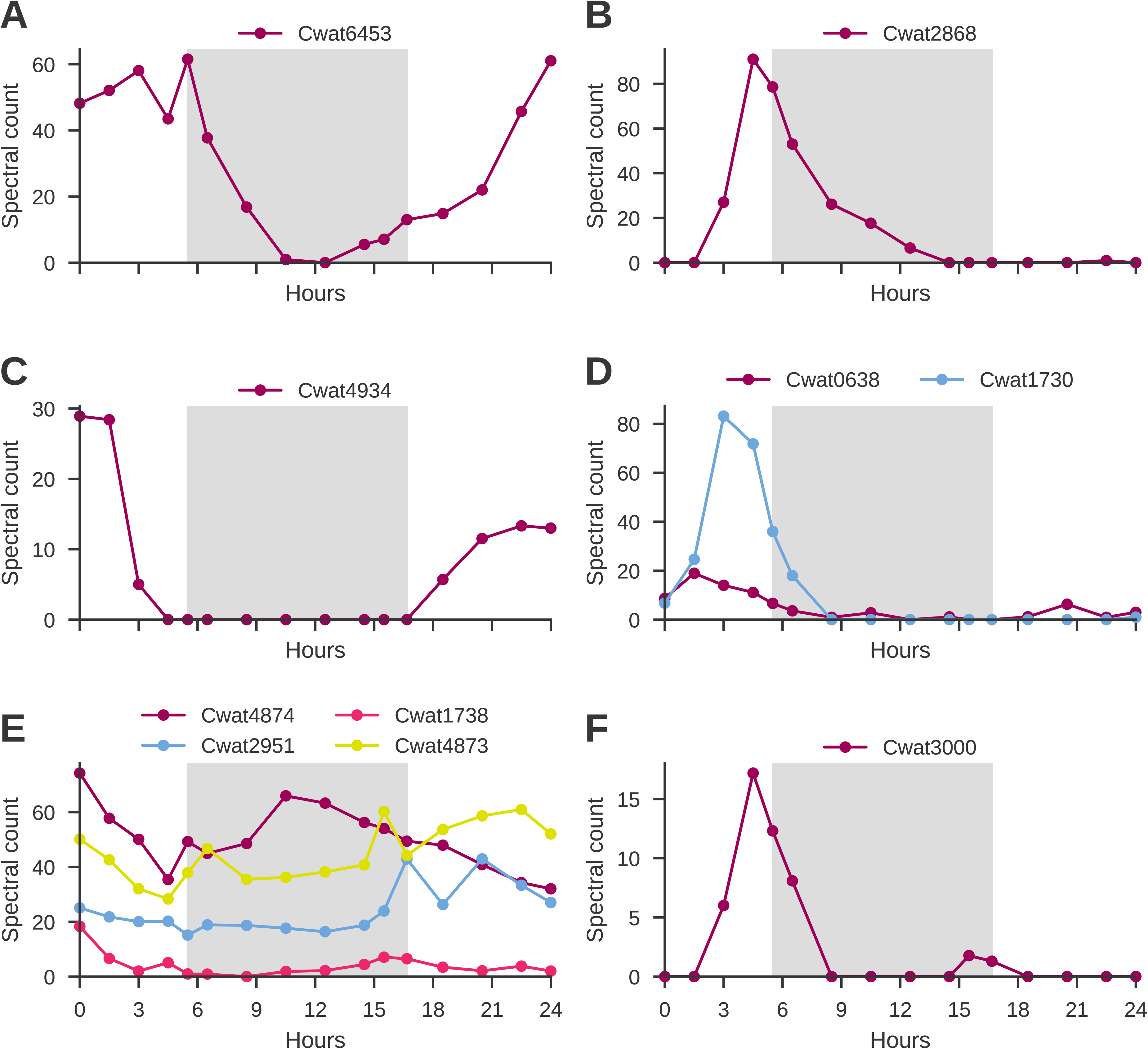
Temporal variability of selected proteins. **A.** Abundance of proteins RpaA (Response regulator receiver:Transcriptional regulatory protein, C-terminal; Cwat6453; Q4C9J3); **B.** the putative oxidoreductase Cwat2868 (oxidoreductase activity; FAD/NAD binding domain); **C.** Peptidase S13 (Cwat4934); **D.** Peptidase M48 (Cwat0638 & Cwat1730); **E.** ClpP, Peptidase S14 (Cwat4874, Cwat1738, Cwat2951, Cwat4873); and **F.** Ferrochelatase (Cwat3000).

Mean abundance patterns with marked changes at dawn and dusk are also observed in one of the data driven clusters, cluster 6 (**Fig 2**), and thus represent candidates for changes in protein abundance linked with diel-related physiological changes. It is interesting to note that this cluster is enriched in the GO term ‘oxidoreductase complex’ which is defined as any protein complex that possesses oxidoreductase activity, and this term is otherwise only observed in cluster 1, as described above in the context of nitrogen fixation. The observation of increased machinery for oxidoreductase activity, such as the hypothetical protein Cwat2868 identified by BLASTp as having oxidoreductase activity with a FAD/NAD binding domain (**Fig 6B**), at the transition (L to D) and concentration of light dependent quinones (**Fig 5E**), may be components of environmental cues for entraining the rhythmicity of the circadian clock to environmental stimuli, in support of the proposed oscilloid model. In the model, and indeed as observed across cyanobacterial lineages, there is no direct photoreceptor, and the redox state of the cell (quinones) (**Fig 5E**) is thought to be sensed^59^; however, unlike in *E. coli* for which there are well known redox sensors and other cyanobacteria with homologs to the activator cikA, a histidine kinase or other sensor with this role is yet to be identified in *C. watsonii*.

### Protein turnover

Protein degradation and translation are critical for maintaining a highly dynamic diel cycle and the cellular controls on protein abundance are therefore of great interest. We identified 49 proteins representing 28 distinct peptidase annotations **(Table 3)**. These peptidases differ based on their structure, such as serine vs. metallopeptidase, and by the type of cleavage they perform. Peptidases involved in peptidoglycan remodeling are observed across all six temporal abundance patterns, playing a key role in synthesis and remodeling of the cell wall during the day, such as D-Ala-D-Ala carboxypeptidase C (Peptidase S13; Cwat4934; **Figure 6C**), and perhaps cell permeability during the night. Peptidases with each of the 10 cleavage types observed in the proteome have maximum abundance during the day (cluster 4), exemplifying the breadth of metabolic activities being performed. Endopeptidases, which can play a role in membrane protein regulation, are represented in clusters 1 and 3-6. Signal peptide cleavage peptidases were observed with maximum abundance during the day (clusters 3-5) and in cluster 6 at the day/night interface (**Figure 6D**).

**Table 3.** Peptidases organized by cleavage type, cluster number, and the time of maximum protein abundance for each cluster. Peptidases are labeled by their abbreviation in the M or S categorization, corresponding to a metallopeptidase or a serine peptidase, as applicable. A key to CwatDRAFT numbers can be found in SI File 2. Other abbreviations used: POP = prolyl oligopeptidase; OpdA = Oligopeptidase A; LAP = Leucyl aminopeptidase; SPI = Signal peptidase I; ClpP = Peptidase S14; GT51 = glycosyltransferases in family 51.

|  | **Cluster** |  |  |  |  |  |
| --- | --- | --- | --- | --- | --- | --- |
| **Cleavage Type** | 1 | 2 | 3 | 4 | 5 | 6 |
| Aminopeptidase |  | LAP |  | M1 |  |  |
| Carboxypeptidase |  | M20D |  | M20D |  |  |
| C-terminal processing |  |  |  | S41A(2) |  | S41A |
| Endopeptidase | M61 |  | M16, M41 | M41, OpdA | M41 (3) | M16 (3) |
| General protein processing |  |  | ClpP (2) | ClpP | ClpP, S8/S53 | S1 |
| Peptidoglycan remodeling | M22 | M50 (2) | M23B (2) | S13, family 51 | S51, M15B / M15C | S13, M52 |
| Proline-specific cleavage |  |  |  | S49 | S9, POP |  |
| Regulation of DNA gyrase activity | U62 |  |  | U62 (2) | U62 |  |
| Signal peptide cleavage |  |  | S24 | SPI (2) | M48 | M48 |
| Unknown |  |  |  | Unspecified (1) |  | Unspecified (1) |
|  | Night | Night / Interface | Early day | Early / Late day | Late day | Interface |
|  | **Time of max protein abundance per cluster** |  |  |  |  |  |

In *Synechococcus elongatus* PCC 7942, Clp proteins contribute to proteome remodeling, which is essential for circadian clock function. Four copies of ClpP are observed in the *C. watsonii* proteome throughout the diel cycle; however, their abundance varies temporally. Specifically, two copies exhibit peak abundance in the early day (cluster 3), one in both early and late day (cluster 4), and one predominantly in the late day (cluster 5) (**Table 3**; **Figure 6E**). This temporal distribution suggests a potential regulatory function linked to circadian rhythms. We therefore contemplate whether proteins such as NifU and NifH are still observed during the light period due to incomplete non-specific protease activity, or whether there is sufficient specificity by which the temporal abundance changes in the wide array of peptidases observed maintain a pool of these proteins while fully degrading other components of the nitrogenase enzyme.

We also observe a marked increase in the abundance of ribosomal proteins involved in translation in the hours prior to dark and peaking as the lights are ramped down, with a gradual decrease in abundance until about the midpoint of the dark phase. Ribosomal proteins uS19, S11, L5, S21, L12, uS2, the first three of which are in the same operon, and a YbaB/EbfC family nucleoid-associated protein (Cwat2370) contribute to the significant GO term ‘intracellular non-membrane-bounded organelle’ (corrected p-value of 7.8E-3) in ***cluster 6***.

### Other temporal signatures

Other key light-independent processes are captured by **clusters 1 and 2** (Fig 2). Photosystem II (Cwat3258) is significantly more abundant in dark than light, a trend recapitulated in Saito et al. (2011)^22^ (***cluster 1***). Enrichment of four proteins results in the significantly enriched GO term ‘photosynthesis, dark reaction’ in ***cluster 2***: (i) Light-independent protochlorophyllide reductase subunit B; Gene: chlB (Cwat0967); (ii) Ribulose bisphosphate carboxylase large chain; Gene: cbbL (RuBisCO) (Cwat2714); (iii) Light-independent protochlorophyllide reductase iron-sulfur ATP-binding protein (Cwat4016); and (iv) Light-independent protochlorophyllide reductase subunit N; Gene: chlN (requires ferredoxin; iron-sulfur clusters) (Cwat4018). Increased abundance of these proteins which have light-independent roles in metabolism concurs with the general temporal abundance pattern in cluster 2. These proteins also contribute to the enrichment of the terms ‘carboxysome,’ the cellular compartment for RuBisCO, and ‘thylakoid membrane’ as light-independent processes also occur in the chloroplast, specifically the stroma. Furthermore, we observe the molecular function GO term ‘glycogen (starch) synthase activity’. As a mature reserve polymer, glycogen has been shown to accumulate during the latter half of the diel cycle and used to power nighttime nitrogenase activity in the related Syn. WH8103^60^. Glycogen has been shown to accumulate in *C. watsonii* WH8501 during the day and at dusk, with a dramatic decrease in abundance during nighttime nitrogen fixation, as seen in Saito et al.^22^. However, starch synthase (Cwat6377) does not display diel variability in this study, perhaps indicating no metabolic gain from targeted enzyme degradation or it may be active during much of the diel cycle. There is speculation that glycogen may also play a role in respiratory protection of the cell, acting as a barrier to oxygen during the dark period prior to use in metabolism (dusk).

Across both clusters 1 and 2 which have increased abundances during the night, the terms ‘metal binding’ and ‘metal cluster binding’ as well as ‘4 iron, 4 sulfur cluster binding (Fe_4_S_4_)’ and ‘iron-sulfur cluster binding’ are enriched in the GO analysis. In addition, both include ‘oxidoreductase activity’ which in cluster 2 is acting on the “CH-CH group of donors” with NAD or NADP as acceptor and in cluster 1 is acting on hydrogen, aldehyde or oxo, and iron-sulfur proteins as donors with iron-sulfur protein and dinitrogen as acceptors. In addition to redox processes, we also detect maximum abundance of three vitamin biosynthesis proteins (thiamine, riboflavin, and cobalamin), a copper-translocating ATPase, and ferrochelatase (**Figure 6F**) at these temporal interfaces. Increased protein abundance at these transitions may align with the shifting metabolic demand for vitamin and metal cofactors. For example, ferrochelatase (Cwat3000) catalyzes the insertion of iron into protoporphyrin IX to form heme, a critical cofactor for cytochromes that supports photosynthetic electron transport proteins including respiratory proteins like cytochrome c oxidase at dusk and the cytochrome b6f at dawn (**Figure 6F**), consistent with the distinct large and small peaks observed in ferrochelatase, respectively, immediately prior to the light shift In this manner the ferrochelatase operates in parallel with the iron-sulfur biosynthesis systems described above for nitrogenase, contributing to the critical need to distribute iron atoms to multiple biosynthetic systems at various temporal transitions. However, the specific connections the responsible proteolytic enzyme(s) (Table 3) that induces their limited diel duration remains to be elucidated.

## Discussion

*C. watsonii* WH8501 displays a remarkably dynamic diel cycle when compared to other abundant unicellular cyanobacteria such as *Prochlorococcus* and other non-diazotrophic microorganisms. Restructuring of the proteome is a non-trivial, yet daily, task for the unicellular diazotroph. Due to temporal separation of photosynthesis and nitrogen fixation, one of the greatest challenges that *C. watsonii* faces is preventing oxidative damage to nitrogenase when oxygen is still present in cells after photosynthesis has ceased. A number of observations point towards a multi-faceted role for respiratory protection, as also postulated in modeling studies^17,21,61^. Beyond respiratory protection mechanisms, the sequential translation and ensuing temporally separated maximal abundance of nitrogen fixation proteins plays a key role in enabling biological nitrogen fixation.

While gene expression studies have shown that the nif regulon is expressed with diel variation^10^, we observe that the gene products accumulate with maximal abundance at different time points. In fact, they increase in abundance in a sequential fashion, beginning with cofactor biosynthesis, then the MoFe nitrogenase protein and lastly the nitrogenase iron protein. This choreographed translation is likely due to the variable stability of individual components and the fully active nitrogenase complex in the presence of oxygen, or the step-by-step needs of Nif complex assembly. Given the current interest in collecting (micro)organismic diel data in the field, these high-resolution temporal laboratory data are a valuable resource to help interpret complex field results and to optimize sample collection timing. The collection of protein datasets in the field can then in turn contribute to accurately capturing potential new nitrogen due to nitrogen fixation in global biogeochemical models, which will be increasingly more important given the shifting and likely expanding range of growth for nitrogen fixers such as *C. watsonii*.

## Data availability statement

The mass spectrometry proteomics data have been deposited to the ProteomeXchange Consortium via the PRIDE^62^ partner repository with the dataset identifiers (reviewers can request a reviewer login). Processed data are available in the Supplementary Information.

All code and analysis notebooks to generate the figures and analyses presented in this manuscript are freely available on GitHub at https://github.com/jgauglitz/Croco_diel under the open source Apache 2.0 license.

## Acknowledgements

We would like to thank the National Science Foundation for NSF OCE PRF awarded to JM Gauglitz (1421196). This work was supported by a grant from the Simons Foundation (LS-ECIAMEE-00001549, Inomura), NSF OCE 2227425 to K. Inomura and 2123055, 2125063, 2048774 to M. Saito and NIH R01GM135709, and the Center for Chemical Currencies on a Microbial Planet (NSF 2019589).

A very special thanks to John B. Waterbury for insightful discussions and mentorship and Frederica Valois for her support, scientific guidance, and assistance in the lab. This manuscript is dedicated to you, Freddy.

## Supplementary Materials

**SI File 1** Croco8501_VB_Nov2008.fasta

**SI File 2** → raw data tables as separate tabs

**SI File 3** GO data - GO term outputs as separate tabs - for cluster 1-6; and sub-clusters 1 (zipped folder)

**SI Table 1** VolcanoPlot_Croco_annotations

## Supplementary Figure Captions

**SI Figure 1.** *Crocosphaera watsonii* WH8501 diel growth. Cellular abundance across the sampling time measured by flow cytometry. Note that cellular division in *C. watsonii* WH8501 typically occurs in the light period, as observed.

**SI Figure 2.** Nitrogen fixation rates, estimated by the proxy measurement made by the acetylene reduction assay. Max nitrogen fixation (between 10.5 and 12.5 hrs) coincides with the maximum abundance of the nitrogenase complex proteins, indicating that the complex is assembled and enzymatically active.

**SI Figure 3.** Changes in protein abundance over the diel cycle for the KaiABC proteins of *C. watsonii* WH8501.

